## Supplementary figures and images for "Genetic and environmental drivers of large-scale epigenetic variation in *Thlaspi arvense*"

### S1 Fig.

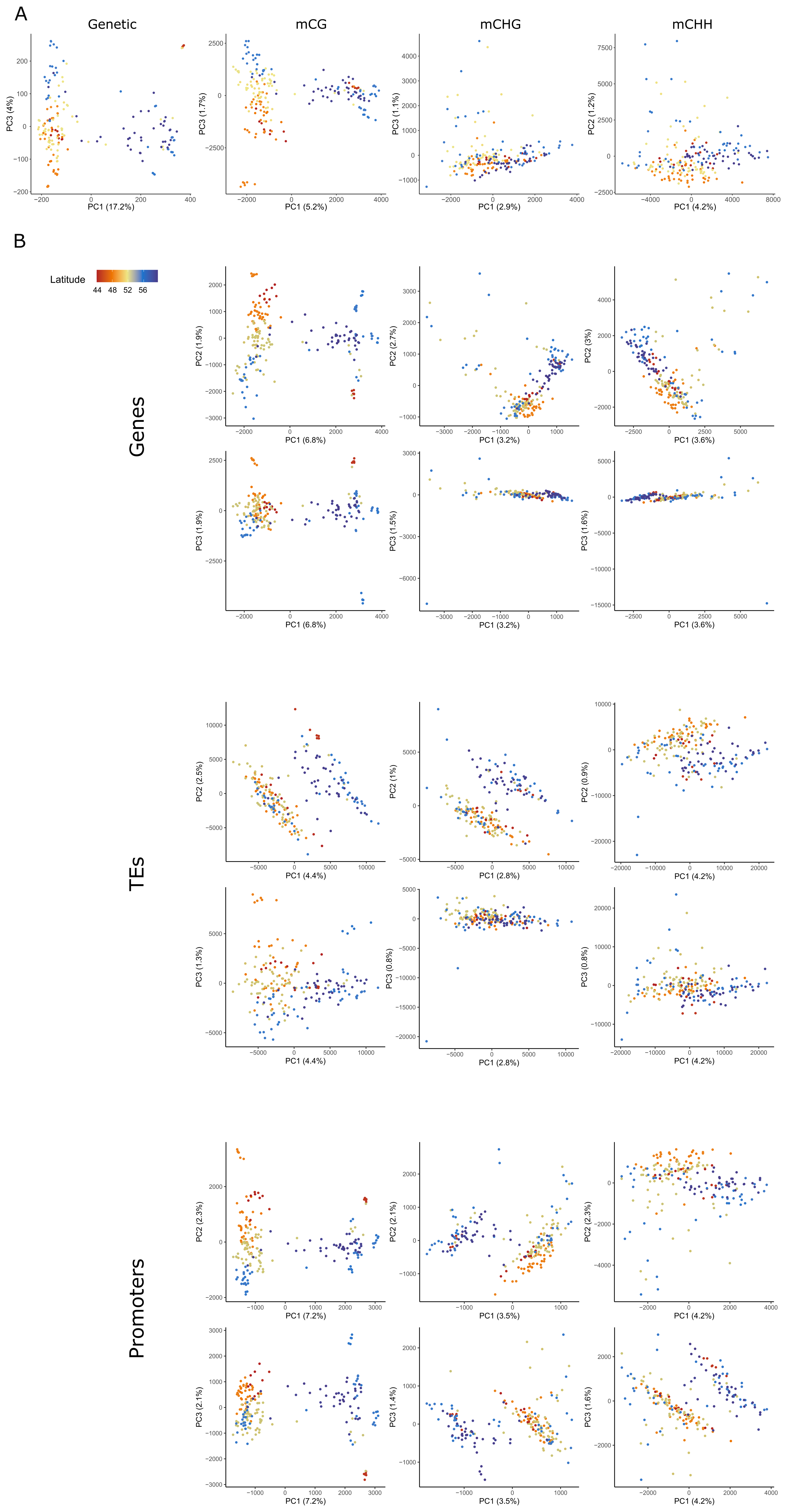

### S2 Fig.

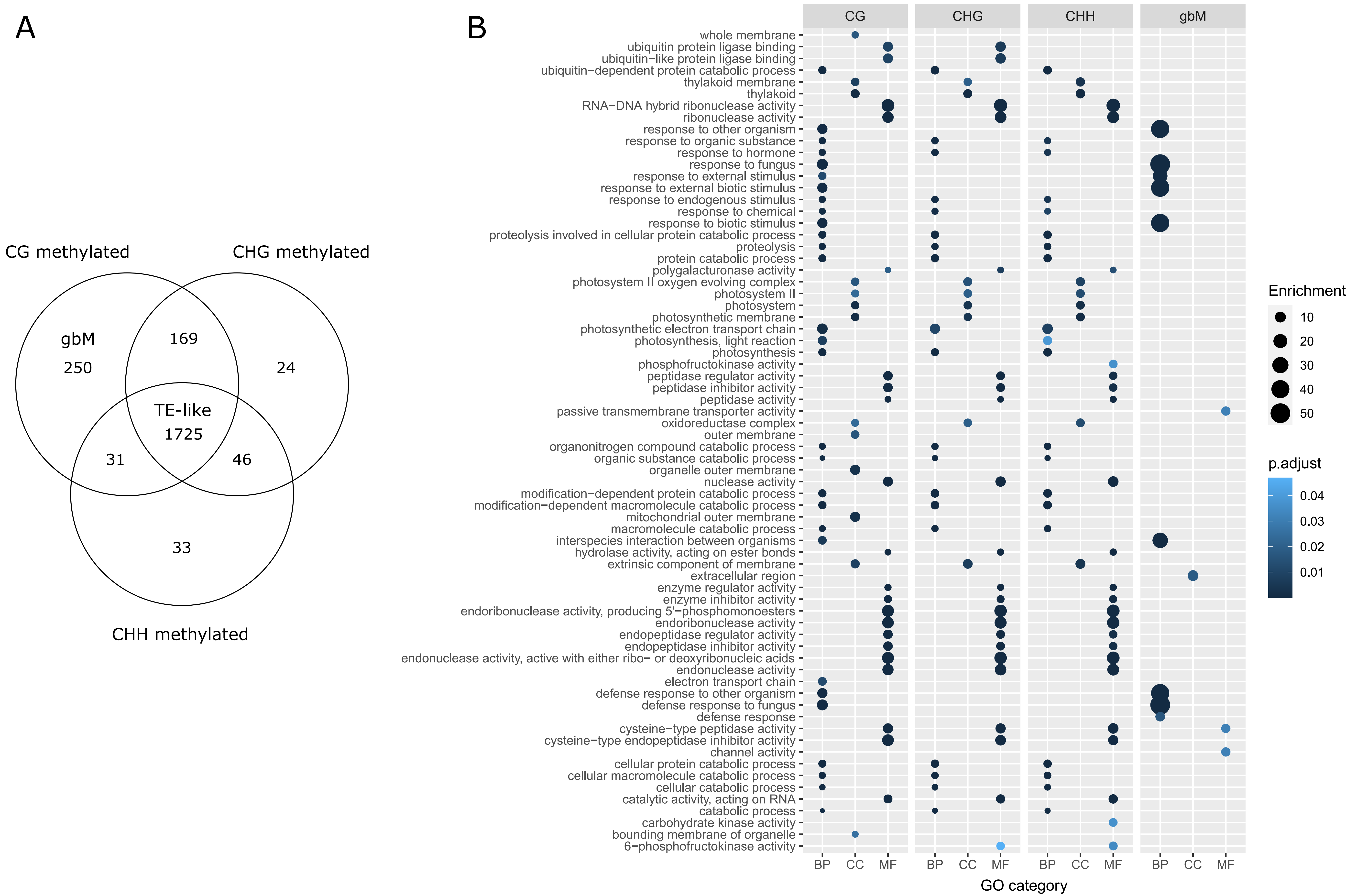

### S3 Fig.

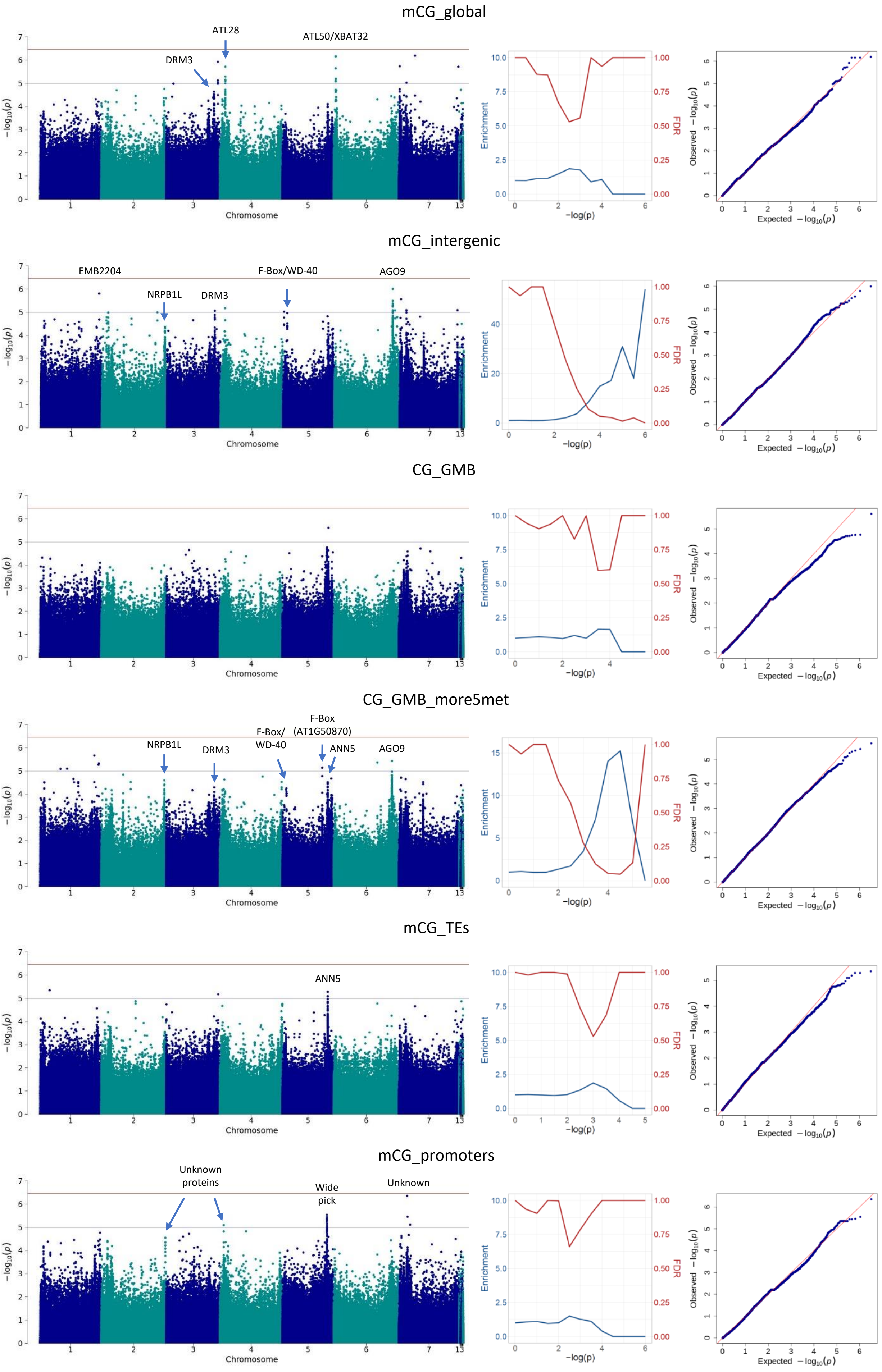

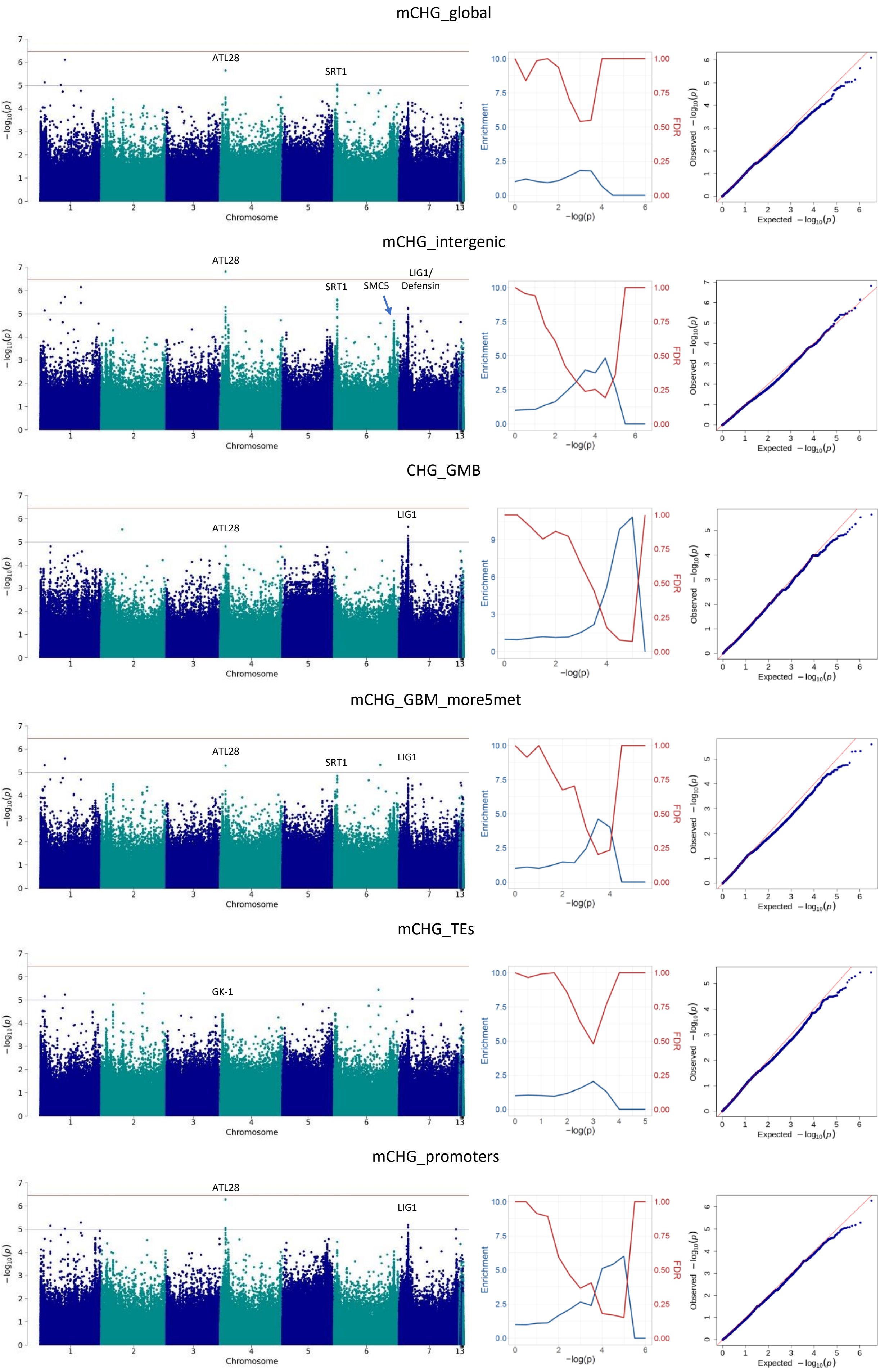

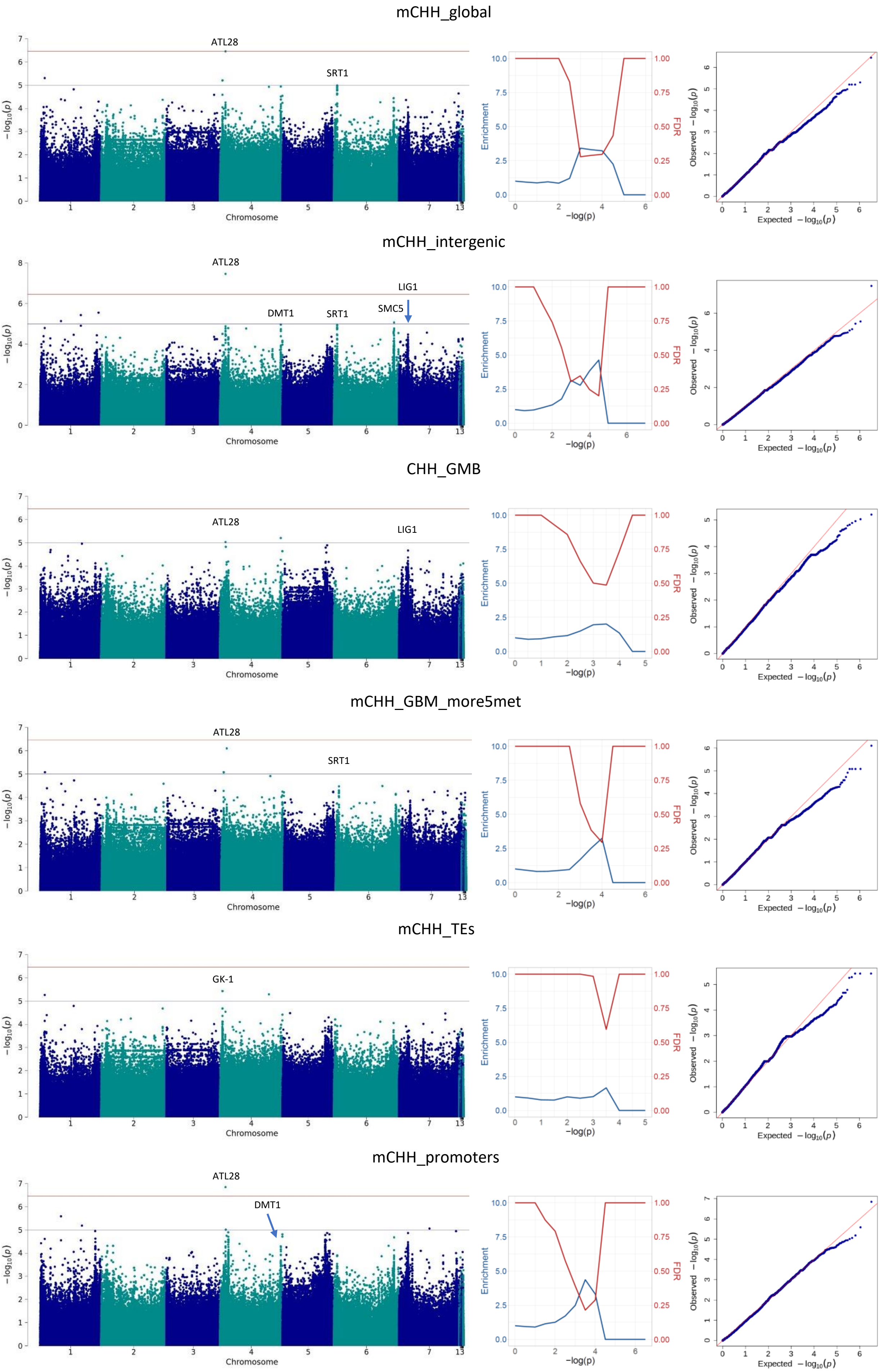

### S4 Fig.

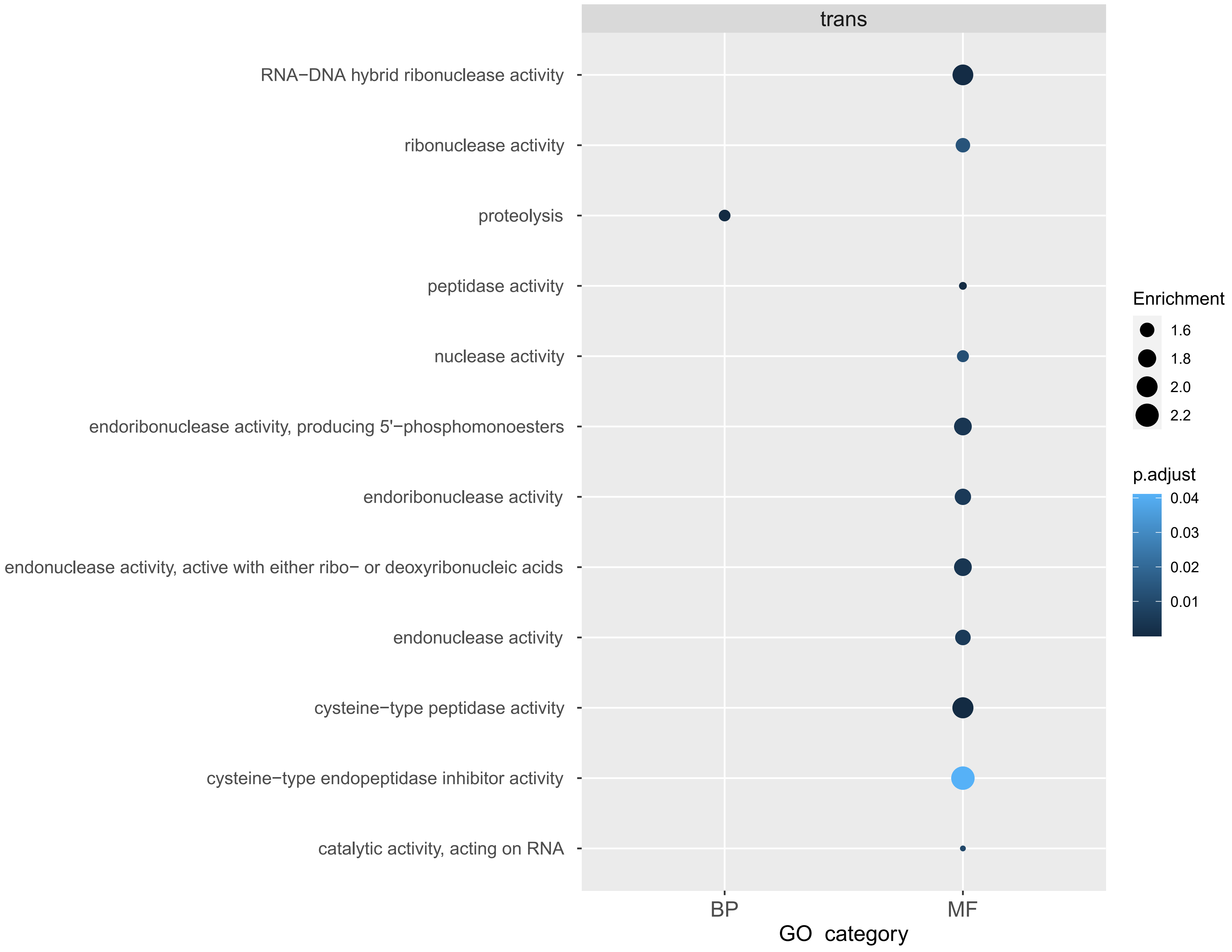
