## Supplementary material for "Genetic and environmental drivers of large-scale epigenetic variation in *Thlaspi arvense*": S1 Table.

**S1 Table. Geographic locations of all *T. arvense* populations.**

| Region | Population | N° lines | Closest town | Latitude | Longitude | Altitude (m) |
|---|---|---|---|---|---|---|
| France | FR_01 | 5 | Les Rives | 43.8508071 | 3.282875 | 743 |
| France | FR_02 | 4 | Mostuéjouls | 44.2385639 | 3.1596174 | 848 |
| France | FR_03 | 6 | Miscon | 44.6281220 | 5.523118 | 821 |
| South Germany | DE_01 | 6 | Tübingen | 48.5402860 | 9.034686 | 458 |
| South Germany | DE_02 | 6 | Löffingen | 47.8766732 | 8.427379 | 708 |
| South Germany | DE_03 | 6 | Braunlingen | 47.9202121 | 8.434509 | 769 |
| South Germany | DE_04 | 6 | Balingen | 48.2802594 | 8.837293 | 539 |
| South Germany | DE_06 | 6 | Hirrlingen | 48.4121117 | 8.8835621 | 431 |
| South Germany | DE_07 | 6 | Empfingen | 48.3843302 | 8.7295661 | 515 |
| South Germany | DE_08 | 4 | Wittershausen | 48.3252288 | 8.6434947 | 536 |
| The Netherlands | NL_01 | 6 | Wageningen | 51.9550616 | 5.6372513 | 8 |
| The Netherlands | NL_02 | 4 | Veenendaal | 52.0403204 | 5.5523601 | 8 |
| The Netherlands | NL_03 | 6 | Herwen | 51.8861228 | 6.1316716 | 10 |
| North Germany | DE_09 | 6 | Halle | 51.5138186 | 11.9201814 | 83 |
| North Germany | DE_10 | 6 | Schwittersdorf | 51.5626351 | 11.7071825 | 187 |
| North Germany | DE_11 | 6 | Eisleben | 51.5409550 | 11.5963349 | 221 |
| North Germany | DE_12 | 6 | Halle | 51.5457637 | 11.9575064 | 90 |
| North Germany | DE_13 | 6 | Plötz | 51.6362233 | 11.9366542 | 82 |
| North Germany | DE_14 | 6 | Rothen | 51.7079562 | 12.0079172 | 88 |
| North Germany | DE_15 | 6 | Bossdorf | 52.0151260 | 12.583979 | 151 |
| North Germany | DE_16 | 6 | Coswig | 51.8852458 | 12.393296 | 55 |
| South Sweden | SE_01 | 6 | Lund | 55.7224293 | 13.184996 | 48 |
| South Sweden | SE_02 | 6 | Lund | 55.7316952 | 13.2524585 | 74 |
| South Sweden | SE_03 | 6 | Lund | 55.7729103 | 13.2571215 | 21 |
| South Sweden | SE_04 | 6 | Eslöv | 55.7483891 | 13.3845241 | 24 |
| South Sweden | SE_05 | 6 | Veberöd | 55.6372295 | 13.5000471 | 34 |
| South Sweden | SE_06 | 6 | Vressel | 55.6676921 | 13.6234294 | 20 |
| South Sweden | SE_07 | 6 | Vanstad | 55.622051 | 13.835959 | 81 |
| South Sweden | SE_08 | 5 | Onslunda | 55.6085961 | 14.0608974 | 108 |
| Central Sweden | SE_09 | 5 | Stockholm | 59.3695897 | 17.9951086 | 9 |
| Central Sweden | SE_10 | 6 | Sollentuna | 59.4340638 | 17.9218504 | 23 |
| Central Sweden | SE_11 | 6 | Rotebro | 59.4773200 | 17.8533700 | 31 |
| Central Sweden | SE_12 | 5 | Uppsala | 59.8191923 | 17.6535739 | 26 |
| Central Sweden | SE_13 | 6 | Uppsala | 59.8447498 | 17.5101196 | 30 |
| Central Sweden | SE_14 | 6 | Uppsala | 59.908915 | 17.599931 | 26 |
| Central Sweden | SE_15 | 6 | Sigtuna | 59.652574 | 17.6884701 | 15 |
