## Supplementary material for "Genetic and environmental drivers of large-scale epigenetic variation in *Thlaspi arvense*": S2 Table.

**S3 Table. List of epiphenotypes, coverage corrections and transformations applied.**

| Trait | N° of outliers removed | Coverage correction | Transformation applied |
|---|---|---|---|
| mCG_global | 4 | yes | none |
| mCHG_global | 3 | yes | none |
| mCHH_global | 0 | yes | none |
| CG_GBM | 4 | yes | INT |
| CHG_GBM | 0 | no | INT |
| CHH_GBM | 0 | no | INT |
| CG_intergenic | 5 | yes | none |
| CHG_intergenic | 0 | yes | gaussianize |
| CHH_intergenic | 0 | yes | gaussianize |
| mCG_TEs | 2 | no | gaussianise |
| mCHG_TEs | 0 | no | none |
| mCHH_TEs | 0 | no | none |
| mCG_prom | 5 | yes | none |
| mCHG_prom | 1 | yes | INT |
| mCHH_prom | 1 | no | INT |
| CG_GBM_more5met | 4 | yes | none |
| CHG_GBM_more5met | 0 | yes | none |
| CHH_GBM_more5met | 0 | yes | none |
